# The roles of divergent and parallel selection in Amazonian and Andean bird responses to glacial cycles

**DOI:** 10.1101/2025.06.21.660863

**Authors:** Vanessa E. Luzuriaga-Aveiga, Matt J. Thorstensen, Jill E. Jankowski, Jason T. Weir

**Affiliations:** Department of Ecology and Evolutionary Biology, University of Toronto, Toronto, ON, M5S 3B2, Canada; Department of Biological Sciences, University of Toronto Scarborough, Toronto, ON, M1C 1A4, Canada; Biodiversity Research Centre & Department of Zoology, University of British Columbia, Vancouver, BC, V6T 1Z4E, Canada

**Keywords:** Climatic cycle, species pair, Pleistocene, speciation, genomics, differentiation, richness, evolution

## Abstract

Both divergent and parallel selection can contribute to evolutionary change, yet their relative contributions to adaptation and speciation are poorly understood, especially during environmental change. We quantified selective sweep signatures and divergence times in Amazonian avian sister species pairs that either differ in elevation and are expected to have experienced strong divergent selection; or occur in similar lowland habitats and are expected to have experienced strong parallel selection. We found that elevational differentiation resulted in a greater accumulation of selective sweep signatures over the five-million-year timespan covered by our dataset, supporting the importance of divergent selection. Nevertheless, lowland-restricted species pairs accumulated more selective sweeps over the most recent two million years, suggesting that parallel selection, driven by major Pleistocene glacial cycles, produced faster evolutionary change than divergent selection in elevationally differentiated species pairs. Our results highlight parallel selection as an important driver of adaptation and evolution, especially during environmental instability.

## Introduction

Natural selection leading to adaptation is a fundamental force shaping genomic differentiation and biological diversity [1–5]. This genomic change may ultimately lead to the formation of new species due to the build-up of genetic incompatibilities that arise as closely related species differentially adapt [6,7]. Divergent natural selection, driven by environmental differences, can fast-track the formation of reproductive barriers between populations – a process often referred to as ecological speciation [8–10]. Populations can also differentiate under parallel selective forces when exposed to non-divergent environmental changes [11]. In this case, each population adapts to the same environmental changes but in different ways. This results in genomic differentiation that may lead to speciation, especially if adaptations are disrupted in hybrid individuals, in a process often referred to as mutation order speciation [12,13]. Despite divergent selection being a more studied and commonly proposed driver of trait evolution, recent comparative analyses across thousands of terrestrial vertebrate species suggested divergent selection drove differentiation in key ecological traits like body size and bill shape in only 10% of species [14]. These inconsistent findings demonstrate fundamental uncertainties about the roles of divergent versus parallel selection in adaptation and speciation.

The Amazon basin provides an ideal laboratory to investigate the roles of both divergent and parallel selection in evolution. Spanning more than six million square kilometers with an elevation range from near sea level to over six thousand meters along the eastern slope of the Andes, the Amazon basin harbors an extensive array of habitat types. Lowland rainforests are followed by the epiphyte encrusted subtropical montane forest zone at 1,300 to 2,400 meters, above which the temperate montane forest zone occurs to treeline (about 3,300 m), with paramo and puna grasslands above treeline [15] (Fig. 1). Many environmental factors vary with elevation, including both climatic variables (e.g., precipitation, temperature, solar radiation, wind speed, oxygen pressure, atmospheric CO_2_, etc.) and biotic factors (e.g., niche breadth, forest height, foraging strata, species diversity, competition strength, etc.) [16]. Moreover, glacial cycles influenced temperature and possibly rainfall across the Amazon basin. Elevational zones of the Andes shifted downslope during glacial periods. This environmental change over the past several million years was most heavily influenced by the mid-to-late Pleistocene glacial cycles [17–19]. However, unlike in northern latitudes, only the highest elevations of the Amazon basin in the high Andes were directly glaciated. While most Andean species may have simply tracked their preferred environmental conditions by moving downslope and upslope during and after glacial periods, lowland species, often surrounded by river barriers on multiple sides, had nowhere to migrate. Lowland populations may have been forced to adapt *in situ* to changing conditions.

**Figure 1.**
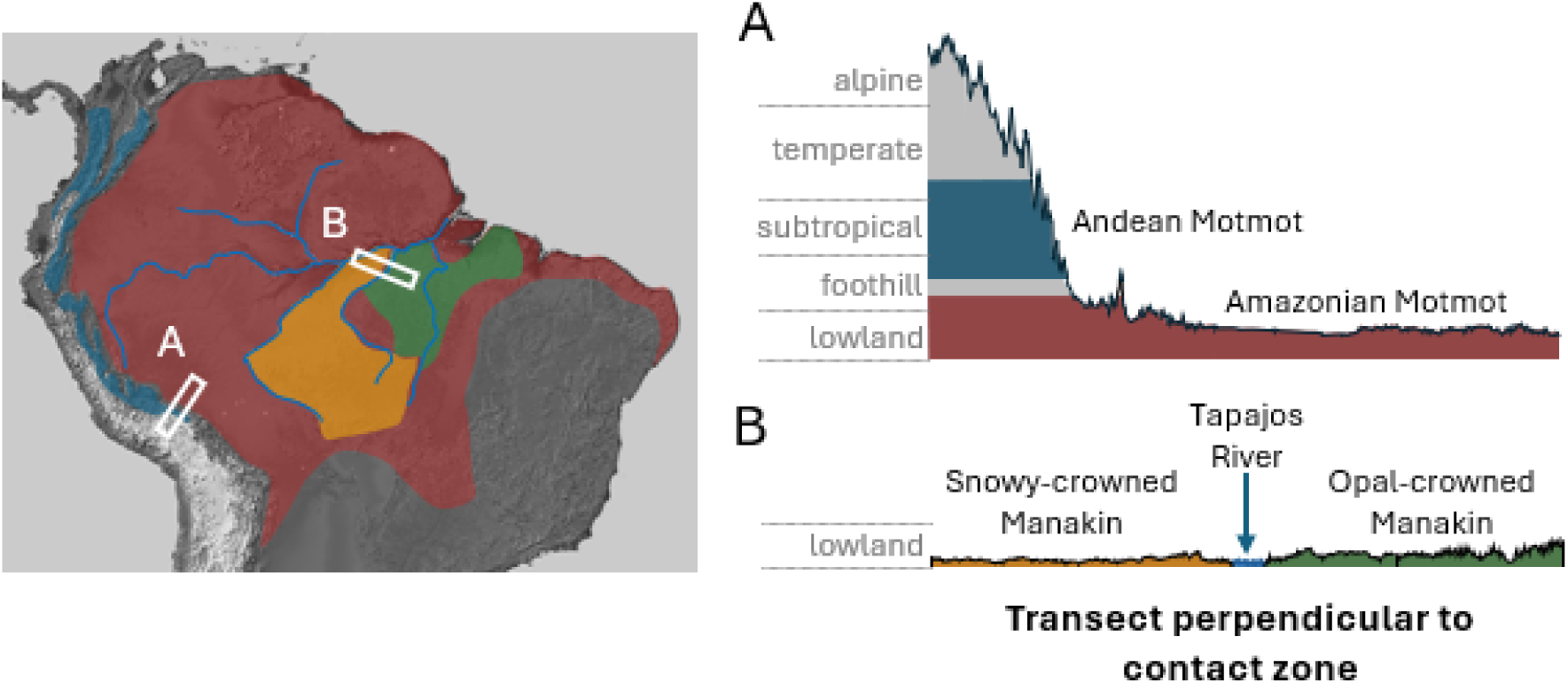
Geographic ranges of two example species pairs overlaid on a topographical map of South America. In A) the foothill forest zone occurs from approximately 600 to 1,300 m (foothill), the epiphyte encrusted subtropical montane forest zone from approximately 1,300 to 2,400 m (subtropical), the temperate montane forest zone to the treeline at approximately 3,300 m (temperate), and the alpine zone exists above 3,300 m. One species pair has the Amazonian Motmot (*Momotus momota*), in red, distributed in lowland forest from 0 to 700 m and its sister species, the Andean Motmot (*M. aequatorialis*), in blue, distributed in Andean cloud forest from 1000 to 2300 m. The other species pair has both species distributed in low elevation rainforest in different regions of the Amazon basin, with the Opal-crowed Manakin *(Lepidothrix iris*) distributed east of the Tapajos River and the Snowy-crowed Manakin (*L. nattereri*) distributed west of the river. Insets A and B show 250 km-long transects running perpendicular to the contact zones between each species pair. Coloured bands show the elevational profiles of each species. Shaded relief topographical base map obtained from NASA (https://www.nasa.gov). Major rivers of the Amazon basin are shown in blue.

These historical patterns of tracking elevation or adapting *in situ* may be evident in the population-level genomic variation of recently-diverged sister species pairs. Specifically, species pairs from different elevational zones would likely experience very different environmental pressures over time as species would first have had to adapt to different environments, then the lowland member of a pair would have continued to adapt to changing conditions over time while the highland member of a pair may have tracked its preferred habitat by migrating up- and down-slope during periods of warming or cooling. Conversely, species pairs with both members from the same elevational zone would have been lore likely to have experienced parallel environmental pressures during glacial cycles resulting in parallel selection as both species in a pair adapted to the same changes in climate [11]. Either circumstance of shared or different elevational zones between members of a sister species pair would have led to an accumulation of selective sweeps after divergence [20], but the selective forces underlying such sweeps could be either divergent or parallel. Therefore, we used species pairs with both members from the lowlands to study parallel selection (acknowledging that some divergent selection may also occur), and species pairs with members from the lowlands and highlands to study divergent selection.

Here, we harness the power of genome-wide data to assess adaptation in closely related pairs of avian species. Positive selection may produce selective sweeps in the genome where the target of selection, and linked nearby regions, become fixed or highly differentiated between populations (high fixation index, *F*_ST_), while genetic diversity within populations (nucleotide polymorphism, π) in these regions is reduced [21–23]. Genomic signatures of strong selective sweeps often stand out as ‘islands of differentiation’ in *F*_ST_ and ‘valleys of genetic diversity’ in π, relative to the genomic background signature of these metrics [20]. We calculated the number of such regions in the genomes for a series of closely related species or genetically differentiated populations within a species (“species pair” hereafter) (Fig. 2). To test the effect of elevational differentiation on the accumulation of selective sweeps through time, we calculated divergence time using site frequency spectrum-based demographic modelling [24]. Species pairs were divided into two categories: lowland-restricted pairs, in which both species in a pair occur in the lowlands of the Amazon (i.e., generally below 1000 m; *n*=9 pairs); and elevationally-differentiated pairs, where the two species in each pair are differentiated in the elevational zones they occupy (*n*=8 pairs). Bayesian linear models were used to assess differences in the accumulation of selective sweeps with respect to divergence time, between the lowland-restricted and elevationally-differentiated species pairs [25].

**Figure 2.**
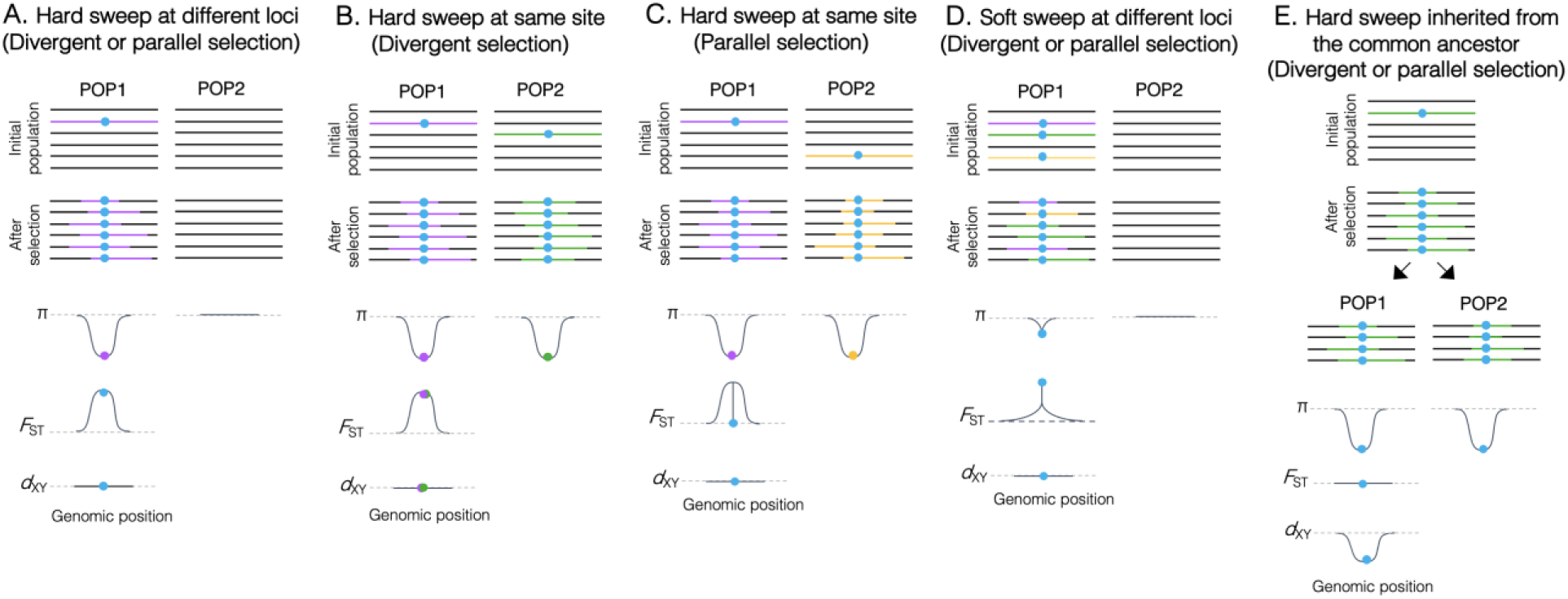
Hard and soft selective sweeps, with corresponding expectations of π, *F*_ST_ and *d*_XY_ under divergent and parallel selection. In hard sweeps (A–C, E), a positively selected de novo mutation (turquoise circle) arises in a single genomic background. Selection causes fixation of this mutation together with closely linked surrounding blocks of the genomic background that have not had time to experience recombination (coloured segments). The soft sweep (D) is similar, but the allele that is the target of selection is segregating in the population prior to becoming positively selected, resulting in all genomic backgrounds it is associated with increasing in frequency. The resulting patterns of π, *F*_ST_ and *d*_XY_ after selection can be used to detect potential selective sweeps. Hard sweeps at different loci (A) result in drops in π only in the species that is the target of selection for a given locus. The drop in π elevates *F*_ST_. Hard sweeps at the same site (B–C) result in a drop in π in both species and an increase in *F*_ST_. They produce almost identical patterns of *F*_ST_ and π under divergent (B) and parallel (C) selection but differ only in the value of *F*_ST_ at the site that is the target of selection, with the site having an *F*_ST_ of 1 under divergent selection (where different alleles at the site of selection are fixed), and an *F*_ST_ of 0 under parallel selection (where the same allele at the site of selection is fixed). Sweeps resulting from divergent or parallel selection cannot easily be discriminated. Soft sweeps at different loci (D) produce similar patterns in π and *F*_ST_ as hard sweeps, but the drops in π and increases in *F*_ST_ are less extreme, making them harder to detect. A sweep in the common ancestor of two populations (E) should result in both daughter populations inheriting low π around the selected site, but, unlike sweeps at the same site in both populations, an ancestral sweep results in a drop in *F*_ST_ and in *d*_XY_ between the daughter populations. Turquoise circles show the allele that is the target of selection. Coloured lines represent different haplotype blocks.

In light of evolutionary theory and natural history of the Amazon basin, we hypothesized that glacial cycles in the mid- to late-Pleistocene drove selective sweeps in birds of the Amazonian lowlands and highlands. More specifically, we hypothesized that the accumulation of selective sweeps between sister species pairs over time would depend on their status as elevationally differentiated or elevationally similar species. We predict that both divergent and parallel selection would lead to an accumulation of selective sweeps over time between members of a species pair. However, divergent selection would lead to a greater rate of selective sweep accumulation than parallel selection as members of a newly diverged species pair adapted to different environments.

## Methods

### Study species

We examined genomic divergence of 17 closely related allopatric species pairs: 9 lowland-restricted pairs, in which both species occur at low elevations in the Amazon basin; and 8 pairs at different elevations, in which one species belongs lower in the Amazonian lowlands than the other in the Andean highlands. In the latter category, both species of the *Chlorospingus* pair are Andean, but one occurring at higher elevations, *C. parvirostris*, than its sister species, *C. flavigularis* (Supplementary Fig. S1). To explore the differences in elevation, we collated elevational information for each species from either a book specializing in Amazonian birds [26] when available, or the Birds of the World online resource if not [27–34]. At least four individuals per species were analyzed (mean = 9.5; range = 4 to 24). Divergence time of species pairs spans from 0.7 Ma to about 6 Ma, representing different stages of the speciation continuum.

Blood or tissue samples of most bird species from the tropical Andes were collected in the Manu Biosphere Reserve in the Department of Cusco or at a variety of locations in the Department of San Martín, Peru. Samples of lowland-restricted species were mostly obtained from Matto Grosso, Brazil. Complementary tissue samples were obtained from Museu Paraense Emilio Goeldi (MPEG), the Louisiana State University Museum of Natural Science (LSUMNS), and the Field Museum of Natural History (FMNH).

### Library preparation and sequence data

A total of 323 individuals were analyzed among 34 species (17 species pairs). DNA was extracted from blood or tissue samples using E.N.Z.A tissue or blood extraction kits (Omega Bio-Tek). Genotyping-By-Sequencing (GBS) libraries [35] were generated for all samples. The Cornell Institute of Genomic Diversity generated libraries for several lowland species in the dataset and these were sequenced on the Illumina HiSeq 2000 platform using 100 bp single-end sequencing. An almost identical GBS method (following the protocol modifications in Luzuriaga-Aveiga *et al.* 2021) was used to generate libraries for the remaining species. The latter were sequenced on the Illumina HiSeq 4000 platform by the Centre for Applied Genomics at the Hospital for Sick Children (SickKids), using 100 bp paired-end sequencing. Sequences of some individuals have already been published by the Weir Lab. Details of individuals used, accession numbers, and sequencing method can be found in the Supporting Online Materials.

Sequences were demultiplexed and filtered using the function *process_radtags* of the STACKS 2.3 pipeline [37], with default parameters. Reads were trimmed for adapters and low-quality sequences using Trimmomatric-0.38 [38], with the following parameters: TRAILING:3 SLIDINGWINDOW:4:10 MINLEN:30. Sequence reads of individuals of each species were aligned to the reference genome of its closest relative available (Supporting Online Materials) using Bowtie2: --phred33 --very-sensitive --mp 5,1 --score-min L, -0.6, -0.6. The *mpileup* and *call* functions of BCFtools 1.12 [39] was used to call genotypes using default settings. Individual genotype calls with < 5x depth and with a genotype likelihood of < 20 were set to missing.

For each species pair we generated three different filtered datasets, each used for different purposes. Dataset 1 was used to calculate *d*_XY_ and π, for which variable and non-variable sites are needed. We excluded sites with more than 90% missing genotype calls and SNPs with high heterozygosity (>0.75). Dataset 2 was used to calculate *F*_ST_, for which only biallelic sites are required. SNPs with a minimum of 4 genotype calls per population were retained. We excluded sites with minor allele frequency <0.05, with mean sequencing depth greater than 95th percentile, and with observed heterozygosity exceeding 0.75. Dataset 3 was used to generate Site Frequency Spectra (SFS). We used identical filtering parameters as in dataset 1, except we removed genotype calls with < 10x depth, and to reduce linkage disequilibrium, we thinned dataset 3 to retain only one SNP per 10 kb.

### Detection of lineage-specific selective sweeps

We first ran a relatedness analysis of individuals within species, using *ngsRelate* [40]. For this analysis, we used unfiltered VCF files of all individuals per species and kept default parameters for the estimation of Jacquard coefficients and relatedness summary statistics. Individuals with co-ancestry coefficient (theta) > 0.1, equivalent to half sibling relationship or higher [41] one of the two related individuals was removed.

Next, we examined signatures of selective sweeps in each of the 17 species pairs, using summary statistics. We used Pixy 1.2.7 [42] to calculate *d*_XY,_ using Dataset 1, and *F*_ST_ and π, using Dataset 2, across 250 kb non-overlapping windows. Only windows with a minimum of 10 SNPs were retained. The female *W*-chromosome was excluded because many individuals were male and we did not have enough female individuals in many comparisons to include the *W*. *F*_ST_, *d*_XY_ and π for each window were transformed into *z*-scores by subtracting the mean across windows and then dividing it by the standard deviation across windows. Mean and standard deviations were calculated separately for each macrochromosome (chromosomes > 2.5-6 μm long [43]; chromosomes 1-10, 1A, 4A and Z) and jointly for the microchromosomes (chromosomes < 2.5 μm long; chromosomes 11-29). One tailed *P*-values were calculated from *z*-scores using the *pnorm* function in R [44], with lower.tail = FALSE to detect significant peaks of *F*_ST_, and lower.tail = TRUE to detect significant drops of π and *d*_XY_. Peaks or drops were considered significant if their *p*-value was ≤ 0.05.

Windows were considered to exhibit signatures consistent with lineage-specific selective sweeps if they possessed significantly elevated peaks in *F*_ST_ and significantly reduced drops in π in one (lineage-specific sweeps) or both (sweeps targeting the same window in both species) species of a pair, and with no significant drop in *d*_XY_. In contrast, a simultaneous drop in *d*_XY_ (Fig. 2D) together with elevated in *F*_ST_ and π in both species of a pair would be consistent with either a selective sweep that occurred in the common ancestor or of a selective sweep that crossed species boundaries through introgression [45]. Selective sweep signatures may span more than one 250 Kb window. To avoid counting these as multiple sweeps, we combined windows of selection less than 1 Mb apart as a single sweep. We standardized the number of detected selective sweeps across species pairs by dividing the number of windows of selection observed by the total number of windows assessed per species pair.

### Estimation of divergence times

We used FASTSIMCOAL2 [24] to estimate the effective population size (*N*_e_) of each member of a species pair, as well as divergence time from a common ancestor. We used Arlequin v3.5 [46] to generate a multidimensional site frequency spectrum (SFS) of species by re-filtering SNPs from down-samples of at least 4 (diploid) individuals per species. We used a minimum site depth cutoff of 10 per individual and manually entered the number of non-variable sites into the SFS. We used family-specific generation times found in the literature to calibrate both per generation mutation rates, calculated from whole-genome datasets in Feng *et al.* (2020) and time of divergence in Myr (Supplementary Table S1). These mutation rates were used to calibrate our demographic models. To build an understanding of the data, we explored a linear model of log_10_*N*_e_ and mean π in a species, with the expectation that the two values should be correlated.

### Patterns of adaptive genomic evolution through time

Using linear regression, we compared how the number of detected selective sweeps accumulated through time for species pairs restricted to the lowlands versus different elevations. If elevational differentiation promotes divergent selection, we would expect to find a higher number of selective sweeps and *y*-intercept (or slope) values in species pairs that occur at different elevations versus lowland-restricted pairs. Potential phylogenetic non-independence of sister pairs was corrected using the following approach. We pruned a time-calibrated phylogeny of all birds of the world [48] to retain only one representative species for each pair in the dataset, using the *keep.tip* function in the R package APE [49]. A phylogenetic multilevel model with the number of detected sweeps dependent on the interaction of divergence time of each pair and category (lowland versus different elevations), and the phylogenetic variance-covariance matrix as a varying intercept was fit in the R package brms [25,50,51]. Specifically, we used a normal distribution to fit the model, with normally distributed priors of mean 0 and standard deviation 0.01 for the model intercept and coefficients, and Student t-distributed priors with 3 degrees of freedom for overall standard deviation and sigma. The model was run over 4 independent Hamiltonian Monte Carlo chains, using 5,000 burn-in and 15,000 sampling iterations each (60,000 iterations total). Adapt delta was raised to 0.999, and maximum tree depth was raised to 12. Trace lots, potential scale reduction factors *R^* of 1, and posterior predictive checks over 100 draws were used to assess model fit. To assess differences between species pairs in the lowlands versus those from different elevations, we used the *hypothesis* function in brms. Specifically, we tested for three differences in brms:

1. To assess whether elevationally differentiated or lowland-restricted species pairs had a greater rate of accumulation of selective sweeps over time, we tested whether the slope of elevationally differentiated species pairs was greater than the slope of lowland-restricted species pairs.
2. To assess the number of selective sweeps at different times due to species pairs being elevationally differentiated or lowland-restricted, we tested whether the number of observed selective sweeps was higher in the birds from different elevations than lowland-restricted birds at 0.7, 2, 3, 4, and 5 Myr. The time at 0.7 Myr was approximately at the start of the mid- to late-Pleistocene glacial cycles.
3. To assess whether there were more selective sweeps in the lowlands-restricted species pairs than the elevationally differentiated species pairs extrapolated to the present day, we tested whether the intercept of the lowland-restricted species pairs was higher than the intercept of the elevationally-differentiated species pairs.

In addition, we used linear modeling to assess the possibility that other patterns may have driven the observed selective sweeps, as opposed to ecologically-driven parallel or divergent selection. Using linear models in R, we tested models of the number of detected sweeps dependent on an interaction of either mean π or log_10_*N*_e_ and elevational category at a species-level, highland or lowland. Another linear model was used to assess whether the accumulation of selective sweeps in highland-lowland pairs was driven by suboscine or oscine species. This was done in a manner consistent with the overall model of selective sweeps and divergence time in brms, except only lowland-highland species pairs were used, and a categorical variable of suboscine or oscine was tested in its interaction with divergence time, rather than elevational category, as previously done. Here, a directional hypothesis test was used to assess whether the suboscine and oscine species each accumulated sweeps over time at a slope greater than 0, as we predicted for the species pairs representing different elevations. We also visualized the proportion of overlap in elevational ranges between lowlands-restricted species pairs and those from different elevations, and plotted those values against the number of observed selective sweeps.

## Results

The species categorized as from highland elevations had generally greater elevational ranges than lowland species (Supplementary Fig. S2). Two species pairs in genera *Chlorospingus* and *Ramphocelus* had substantial overlap in elevational ranges between the low and high elevation species in these pairs, but the populations studied nevertheless appear to be elevational replacements where they co-occur on the same mountain slopes [26].

The standardized number of detected selective sweeps varied between species pairs and elevational categories in a model with overall *R*^2^ of 0.65 (95% credible interval (CI): [0.34, 0.88]) (Fig. 3). Lowland-restricted pairs had a flat relationship between detected selective sweeps and age of the species pair (slope = 8.20×10^-5^, 95% CI: [-1.62×10^-3^, 1.52×10^-3^]) while the pairs from differing elevations had a strong positive increase (slope = 3.14×10^-3^, 95% CI: [7.69×10^-4^, 5.31×10^-3^]). An evidence ratio of 155.66 from a directional hypothesis test corresponded to a 99.4% posterior probability that the pairs from differing elevations had a higher slope than the lowland-restricted pairs. In a test for whether the slope for lowland-restricted birds was > 0, the evidence ratio was 0.83 and posterior probability was 45.5%, thus we could not distinguish this slope from 0. The third and final use of the *hypothesis* function in brms showed that lowland-restricted species pairs had more observed selective sweeps until approximately 2 Myr (evidence ratio 12.09, posterior probability 92.4%), where the number of selective sweeps was indistinguishable between the lowland-restricted and elevationally-differentiated pairs (evidence ratio 1.02, posterior probability 50.4%), while beyond 2 Myr in the past, the elevationally-differentiated pairs had more observed selective sweeps (evidence ratios of 19.48, 91.31, and 143.93 and posterior probabilities of 95.1%, 98.9%, and 99.3% at 3, 4, and 5 Myr respectively). The lowland-restricted species pairs had a higher intercept than the elevationally differentiated species pairs (evidence ratio 23.77, posterior probability 96.0%).

**Figure 3.**
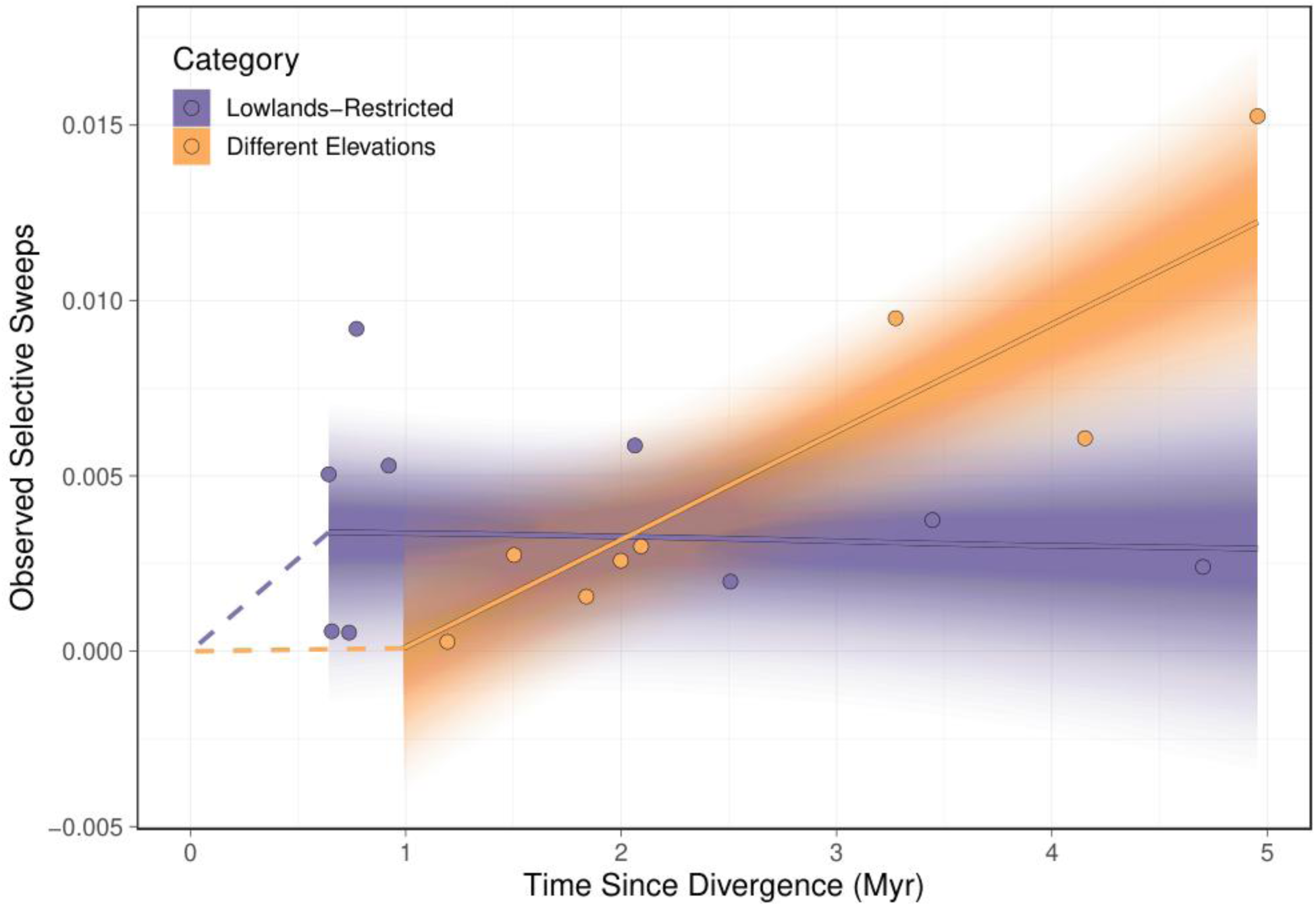
Number of observed selective sweeps accumulated between species pairs versus their divergence times, calculated from the nuclear genome using demographic modeling. Species pairs restricted to the Amazonian lowlands are shown with purple, and pairs occupying different elevational zones from the Amazonian lowlands towards the high Andes are shown in orange. Lines show model fits with phylogenetic relatedness accounted for. The lightest shading shows 95% credible intervals, while darker shading shows 10% credible intervals to emphasize posterior distributions underlying estimates of this linear model. Dashed lines reach from the most recent estimate of each line to the present day of 0 Myr, to emphasize selective sweep accumulation in the mid- to late-Pleistocene.

Observed π or *N*_e_ were correlated (r=0.72, adjusted *R*^2^=0.50, *p*=2.01×10^-6^; Supplementary Fig. S3), and neither drove observations of selective sweeps, (*p*=0.17 and 0.40, respectively) (Supplementary Figs. S4; S5). A paired t-test of pairs at different elevations shows that highland species have consistently lower π but not log_10_*N*_e_ relative to their lowland close relative (paired t-test: *p* = 0.025 (π), *p* = 0.28 (*N*_e_); Fig. 4).

**Figure 4.**
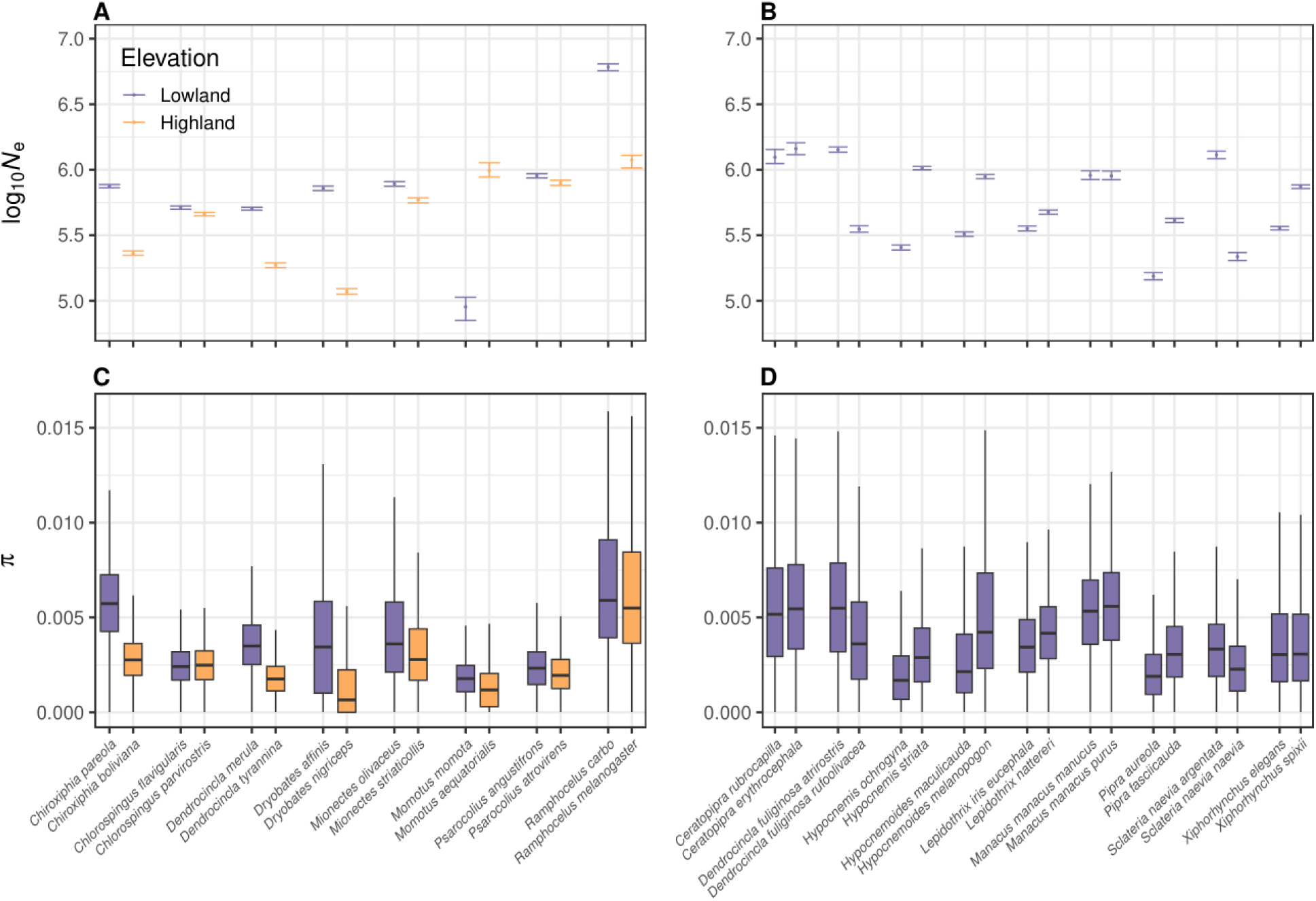
Estimated effective population size (log_10_*N*_e_, top panels A and B) and box and whisker plots of average genome-wide nucleotide diversity (mean π, bottom panels C and D) across 250 Kb genomic windows. Species delineated by category, between pairs from different elevations (A and C) and lowland-restricted pairs (B and D). Lowland species are in purple, while highland species are in orange.

A limitation of our study design is the taxonomic mismatch between species categories. All species pairs from the Amazonian lowlands are suboscines, whereas species pairs at different elevations are phylogenetically more diverse, including two non-passerine pairs (motmots and woodpeckers), three oscines (tanagers and icterids) and three suboscines (woodcreepers, manakins, and flycatchers). However, we found that non-suboscine species in elevationally differentiated pairs accumulated selective sweeps at a similar pace through time as suboscine species, and both exhibited signatures of divergent selection as slopes were greater than 0 (oscine evidence ratio 44.6, 97.8% posterior probability; suboscine evidence ratio 381.2, 99.7% posterior probability) (Fig. S5). Moreover, all linear models were corrected for phylogenetic structure. As such, we do not believe that the differences in the patterns found here between lowland-restricted species pairs and pairs with elevation differentiation were driven by phylogenetic composition.

In addition, no relationship was evident between the proportion of elevational range overlap and the number of selective sweeps in either category of lowlands-restricted species pairs or those from different elevations (Supplementary Fig. S7).

## Discussion

Divergent selection is celebrated as a key driver of rapid evolutionary change, while the contribution of parallel selection is less well understood. Both have the potential to drive a rapid accumulation of selective sweeps. Here we quantified the number of selective sweep signatures accumulated through evolutionary time in related pairs of Amazonian species that differ greatly in elevation and therefore likely experienced stronger divergent selection. We compared these with pairs restricted to similar habitats at low elevations, which likely experienced weaker divergent selection but are expected to have been strongly influenced by parallel selection pressures. We found a substantially steeper increase in the number of selective sweeps through time in elevationally differentiated pairs of species while species pairs at the same elevation showed a flat relationship between sister species age and the number of observed selective sweeps. This difference in slope was robust to phylogenetic structure, independent of genome-wide π or *N*_e_, and is consistent with the long-term effect of ecologically driven divergent selection accelerating evolution over the course of millions of years in elevationally differentiated pairs compared to lowland restricted pairs. Nevertheless, lowland restricted pairs accumulated more selective sweeps over the glacial cycles of the past two million years, which we suggest reflects the power of parallel selection to generate bursts of adaptation and genomic evolution during periods of intense climate change.

For species pairs restricted to lowland regions, selective sweeps did not accumulate as a function of time like they did in elevationally differentiated pairs. This observation would make sense if the major climatic cycles of the past 700 kyr were overwhelmingly responsible for selection compared to earlier periods. Under this scenario all species pairs from the same lowland elevations older than 700 kyr (i.e., all species pairs in our dataset) should have experienced roughly the same amount of selection, regardless of their differing time of divergence from a common ancestor. These climatic cycles are believed to have caused a general cooling and possibly decreased rainfall in lowland Amazonia [52,53]. Unlike highland species which can track their preferred environments downslope during glacial periods, low elevation species have nowhere to go, and are thus forced to adapt in parallel. Mid- to late-Pleistocene climatic cycles would have thus intensified parallel selection pressures to adapt in lowland species while the much more stable environment prior to these cycles would exert much weaker selection. The flat relationship of our lowland restricted pairs between ∼700 kyr and 5 Myr is thus consistent with almost no detectable effect of selection prior to the glacial cycles, but a substantial effect during them from ∼700 kyr to the present. This contrasts with elevationally differentiated pairs of species where the accumulation of selective sweeps through time is strongly positive, suggesting ongoing divergent selection up to 5 Myr was driven by elevational differentiation rather than just the effect of more recent Pleistocene glacial cycles.

In addition to the rate of selective sweep accumulation, the absolute number of observed sweeps at different times was also informative for elucidating the roles for different types of selection. There were more observed selective sweeps due to parallel than divergent selection from the present up until approximately 2 Myr, after which there are more selective sweeps attributable to divergent selection (Fig. 3). Therefore, while divergent selection had a consistent effect over time and led to more selective sweeps in species pairs from different elevations, parallel selection had a more prominent role in evolution during glacial cycles. This interpretation relies on our assumption that species pairs at the same elevations experience mostly parallel selection given the similar environments they are in. While environment is much less variable across lowland Amazonia than it is along the elevational transects of the Andes, some divergent selection driven by fine-scale spatial heterogeneity may have contributed to selective sweeps in lowland-restricted species pairs [54] and our estimates cannot untangle the contribution of these from the effect of parallel selection. Nevertheless, this finding challenges a widespread belief that ecological and environmental differences accelerate speciation (i.e., ecological speciation) and are its key drivers, while parallel selection plays a less important role in speciation [10]. Instead, our results suggest that parallel selection may be capable of driving even faster evolutionary change when environments are unstable and species are forced to adapt *in situ*.

Our methods are unlikely to capture all selective sweeps. First, adaptive alleles do not always generate *F*_ST_ peaks when traits under selection are highly polygenic, because polygenic traits may evolve with incomplete sweeps at many loci. In such cases, the influence of selection on individual loci is likely too small to be detected [23]. Second, soft sweeps acting on standing variation may generate too weak of a signature for our methods to detect, because with standing variation the target of selection will have recombined with a variety of surrounding genetic variants, which then fail to materialize as a sweep signature when the target is fixed (Fig. 2). It thus remains possible that lowland-restricted species did experience selection before the mid- to late-Pleistocene glacial cycles in primarily polygenic traits or soft sweeps. The stronger parallel selection forces induced by the glacial cycles would then have driven an increase in hard sweep signatures more recently which is the primary signal we detect. This scenario is also plausible because greater nucleotide diversity observed in the lowland species (Supplementary Fig. S2) means there is a larger pool of standing genetic variants present that soft selection could have acted upon. Given these considerations, we can only conclude that our different patterns uncovered between species pairs that do and do not have elevational differentiation primarily reflect the effect of hard selective sweeps while the effect of soft sweeps and selection on highly polygenic traits remains unknown.

Another limitation of this work is that genomic architecture may have influenced the results. We analyzed *F*_ST_ and π in windows of at least 250 kb, thus, smaller sweeps may go undetected and their detection will require whole genome sequencing. The larger sweeps we were able to detect may be associated with architecture such as chromosomal inversions, which are understood to support local adaptation and selective sweeps in a range of species [55,56].

Also, centromeres and telomeres possess low recombination rates that facilitate sweeps, which lower π and elevate *F*_ST_ over broad genomic regions, thereby generating strong sweep signatures that are easily detected (see Cutter & Payseur 2013). However, there are only a limited number of these genomic features, and if they were a major contributor to our detections of selective sweeps, the number of such selective sweep signatures we detected might accumulate rapidly over short time scales and then remain constant with increasing time, as sweeps in older species pairs would be overwritten by subsequent sweeps. In such a scenario, the elevationally-differentiated pairs should also exhibit a flat relationship between species pair age and the number of selective sweeps, like lowland-restricted pairs. Instead, we uncovered a strong positive slope for elevationally-differentiated pairs (Fig. 3), indicating that avian genomes are capable of accumulating detectable selective sweep signatures through time. Therefore, the approach we adopted for detecting divergent selection might be influenced by, but was not invalidated by, conserved genomic architecture.

In conclusion, despite the limitations of our approach, we found clear differences between elevationally differentiated and lowland restricted species pairs in terms of selective sweep accumulation through time. Over the five-million-year timescale of our dataset, strong divergent selection associated with elevational differentiation resulted in a much greater number of selective sweeps than seen in lowland restricted pairs, which confirms the importance of divergent selection as has previously been emphasized (e.g., Schluter 2009). Nevertheless, our data suggests lowland restricted species pairs accumulated more selective sweeps over the major Pleistocene glacial cycles consistent with climate change forcing them to adapt *in situ*, while highland species could track their preferred environment up and down slope with less need for adaptation. Parallel selection may thus result in limited adaptation when environments are highly constant through time (i.e., prior to major Pleistocene glacial cycles) but may elevate the rate of accumulation of selective sweeps during periods of environmental instability (i.e., during major Pleistocene glacial cycles) when parallel environmental changes force species to adapt. Parallel selection is likely to target different genes in different populations, leading to genomic differentiation and the accumulation of genetic incompatibilities that ultimately are believed to play a key driving force in Amazonian speciation [57,58]. Our findings are consistent with a recent meta-analysis of birds, mammals, reptiles, and other groups which found preliminary evidence that parallel selection was a generally stronger force in shaping trait evolution than divergent natural selection [14]. Together, these results suggest parallel selection may drive adaptation and speciation during periods of intense environmental change in the most species rich region of the planet. While the role of divergent natural selection has been well documented, there remains a great need for studies to address the contribution of parallel selection to adaptation and speciation.

## Acknowledgements

Else Mikkelsen and Jordan Bemmels provided valuable discussions throughout this study. Ellen Nikelski helped improve an earlier draft. Land-owners of Brazil graciously allowed us to sample on their properties. Alexandre Alexio facilitated Brazilian collecting permits (Brazilian National Research Council 4253-1, 40173-1 and 6581-1). Mauricio Ugarte-Lewis facilitated Peruvian collecting permits (RDG N° 158–2017-SERFOR-DGGSPFFS). The following museums provided tissue samples: Museu Paraense Emlio Goeldi, Louisiana Museum of Natural History, and Field Museum of Natural History. Funding was supplied by the National Secretariat of Science, Technology and Innovation (SENESCYT-IFTH) from Ecuador to V.E.L.A., the Eric and Wendy Schmidt AI in Science Postdoctoral Fellowship, a program of Schmidt Sciences, LLC (to M.J.T.), and Natural Sciences and Engineering Research Council of Canada (NSERC) Discovery Grant RGPIN-2012-418294, RGPIN-2019-04260 (to J.E.J.) and National Science Foundation Grant DEB-1120682 (to J.E.J.), and the Natural Sciences and Engineering Research Council of Canada Accelerator Grant (no. 492890; to J.T.W.), Discovery Grants (402013-2011, RGPIN-2016-0653, RGPIN-2022-04817) to J.T.W.

## Supporting Information

**Supplementary Table S1:** Generation times and mutation rates used in coalescent modelling for major families represented in the dataset.

| Family | Genus | Generation time | Mutation rate |
| --- | --- | --- | --- |
| Icteridae | *Psarocolius* | Mean = 1.83 | $2.096 \times 10^{-9}$ |
| Thraupidae | *Ramphocelus* | Mean = 2.86 | $3.98 \times 10^{-9}$ |
| Passerellidae | *Chlorospingus* | Mean = 2.33 | $2.99 \times 10^{-9}$ |
| Thamnophilidae | *Hypocnemoides*<br>*Hypocnemis*<br>*Sclateria* | Mean = 2.44 | $4.14 \times 10^{-9}$ |
| Dendrocolaptidae | *Xiphorhynchus*<br>*Dendrocincla* | Mean = 2.83 | $4.14 \times 10^{-9}$ |
| Tyrannidae | *Mionectes* | Mean = 1.98 | $3.73 \times 10^{-9}$ |
| Pipridae | *Ceratopipra*<br>*Pipra*<br>*Lepidothrix*<br>*Manacus*<br>*Chiroxiphia* | Mean = 3.34 | $6.06 \times 10^{-9}$ |
| Picidae | *Dryobates* | Mean = 2.86 | $6.89 \times 10^{-9}$ |
| Momotidae | *Momotus* | Mean = 2.85 | $5.84 \times 10^{-9}$ |

**Figure S1.**
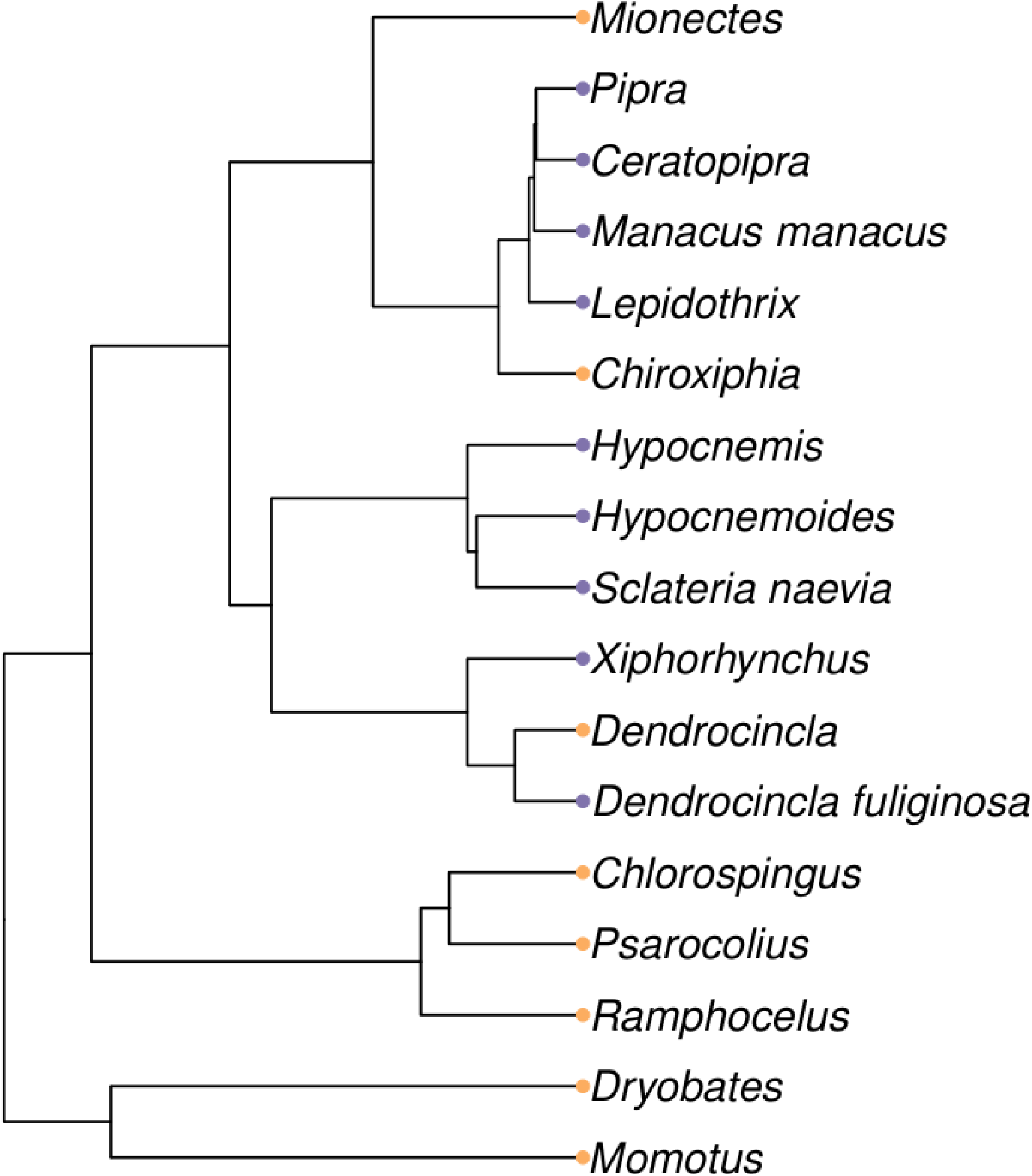
Phylogeny of the taxa included in the dataset. Tips of the tree represent sister species or subspecies pairs, colour-coded by their elevational category: purple for lowland-restricted pairs and orange for pairs at different elevations. *Dendrocincla fuliginosa* from the lowlands represents two subspecies, *D. f. atrirostris* and *D. f. rufoolivacea*. *Dendrocincla* at different elevations represents *D. merula* and *D. tyrannina*.

**Figure S2.**
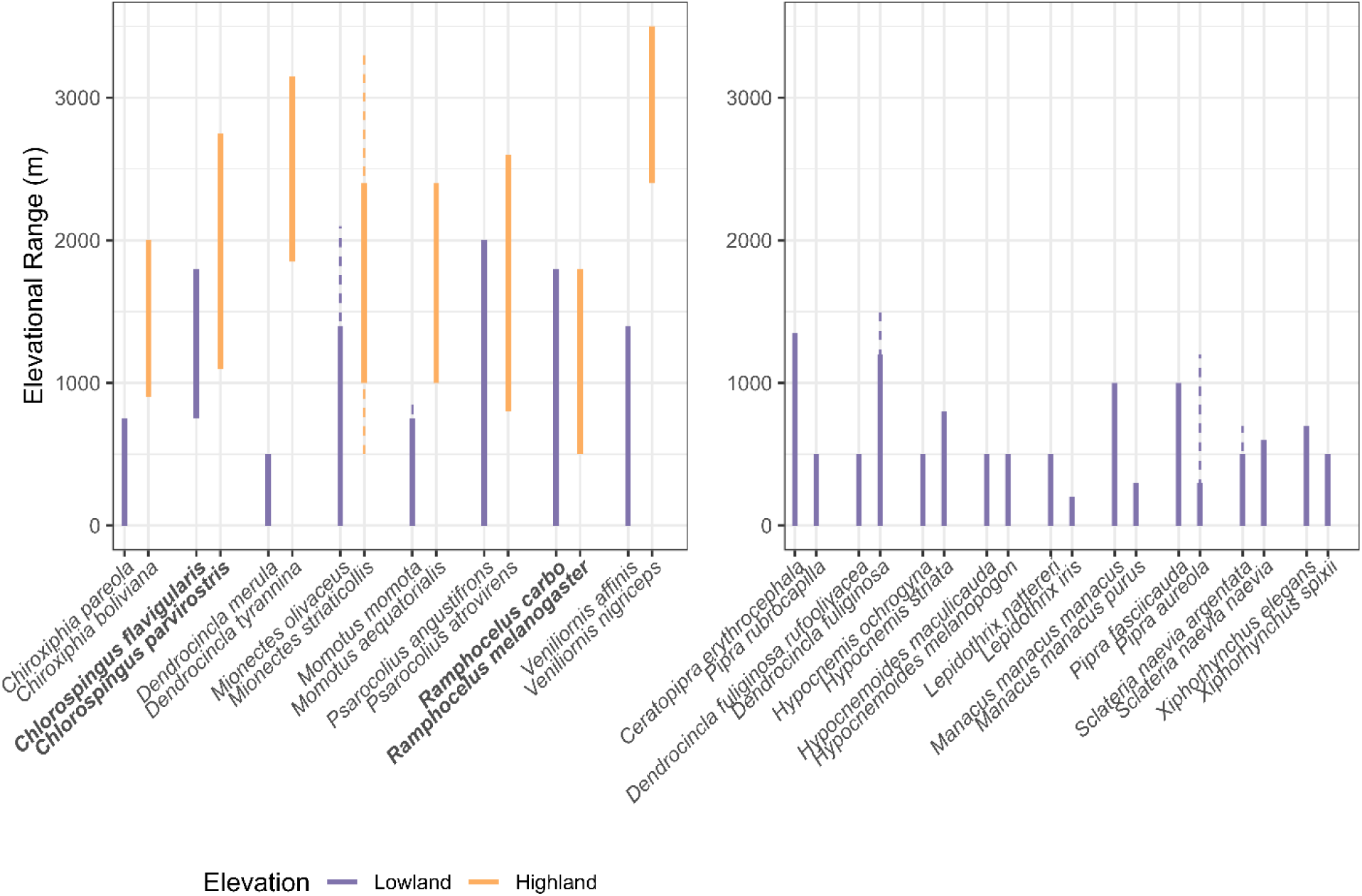
Elevational ranges for species in the present study. For species where both a core range and maximum and minimum observations were recorded, dashed lines were used to show the extreme ends of a range. However, this information was not available consistently. While genera *Chlorospingus* and *Ramphocelus* (bold) had largely overlapping species elevational ranges, they were allopatric and likely experienced different historical environments.

**Figure S3.**
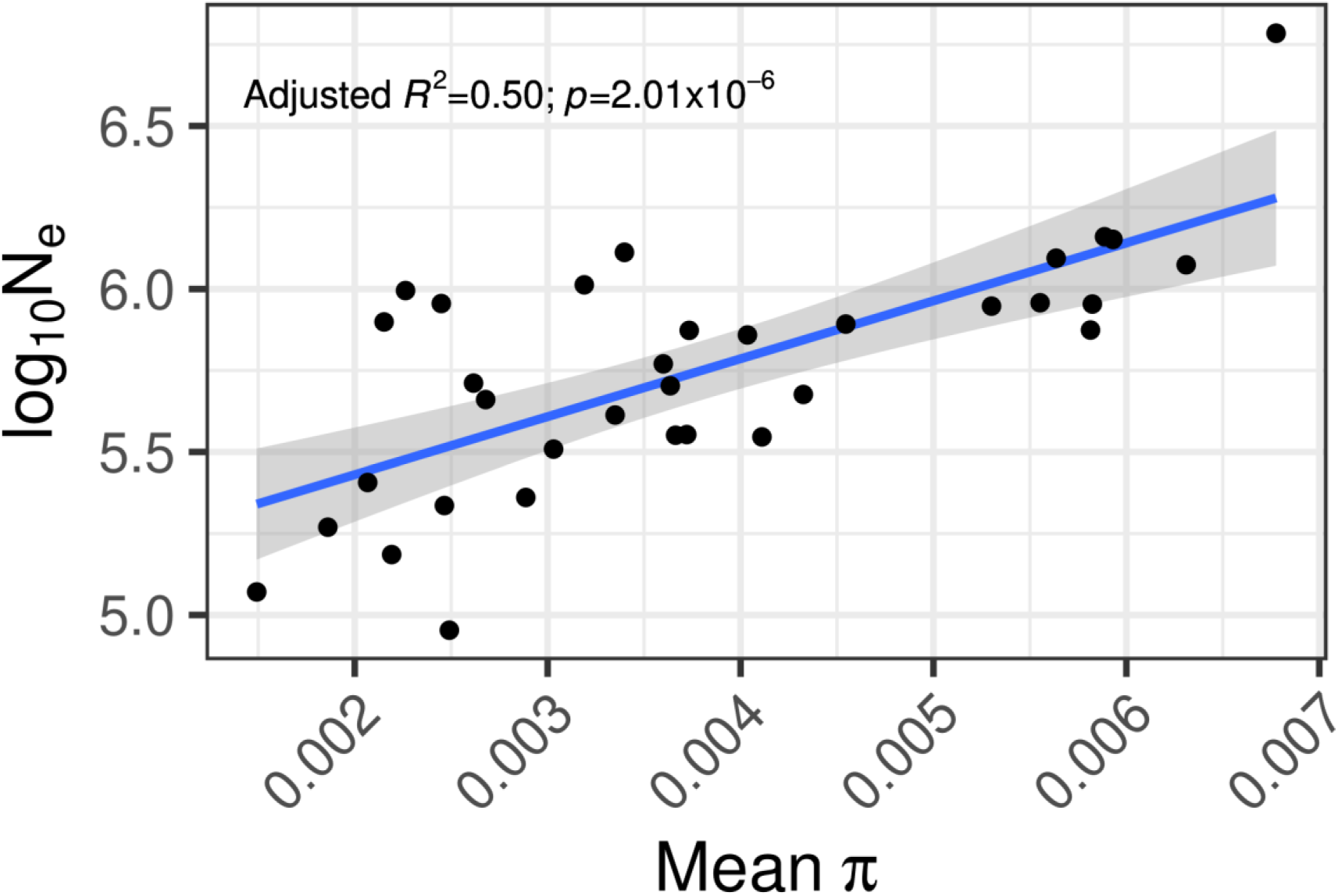
Linear regression between effective population size (*N*_e_) and mean genome-wide nucleotide diversity (π).

**Figure S4.**
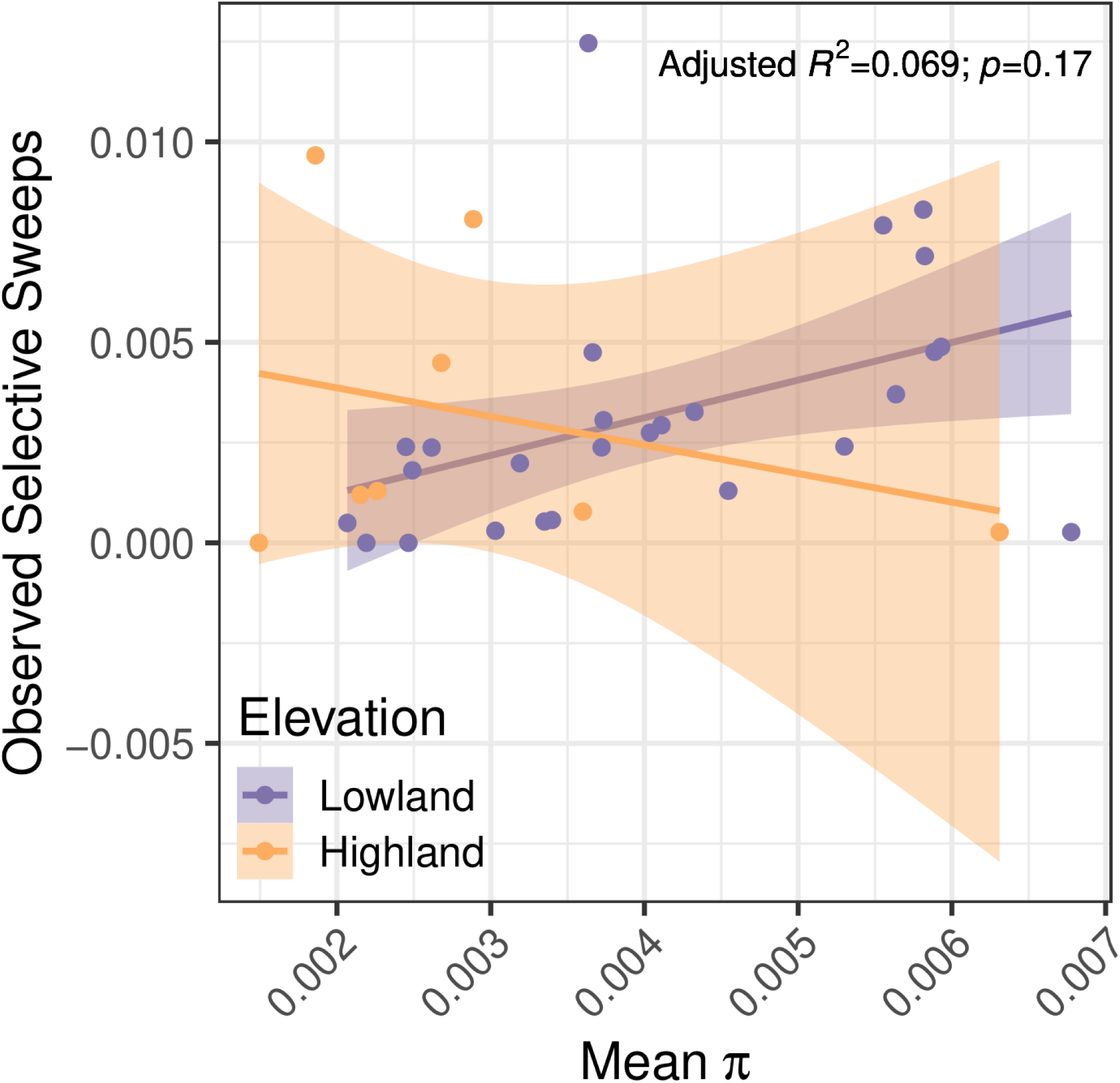
Regression analyses between number of lineage-specific windows of selection and average genome-wide nucleotide diversity, π, for individual species (not species pairs). Here, mean π interacted with elevational category of lowland or highland. Species from the Andean highlands are shown in orange, and lowland Amazonian species are shown in purple. This model was not significant (*p*=0.17).

**Figure S5.**
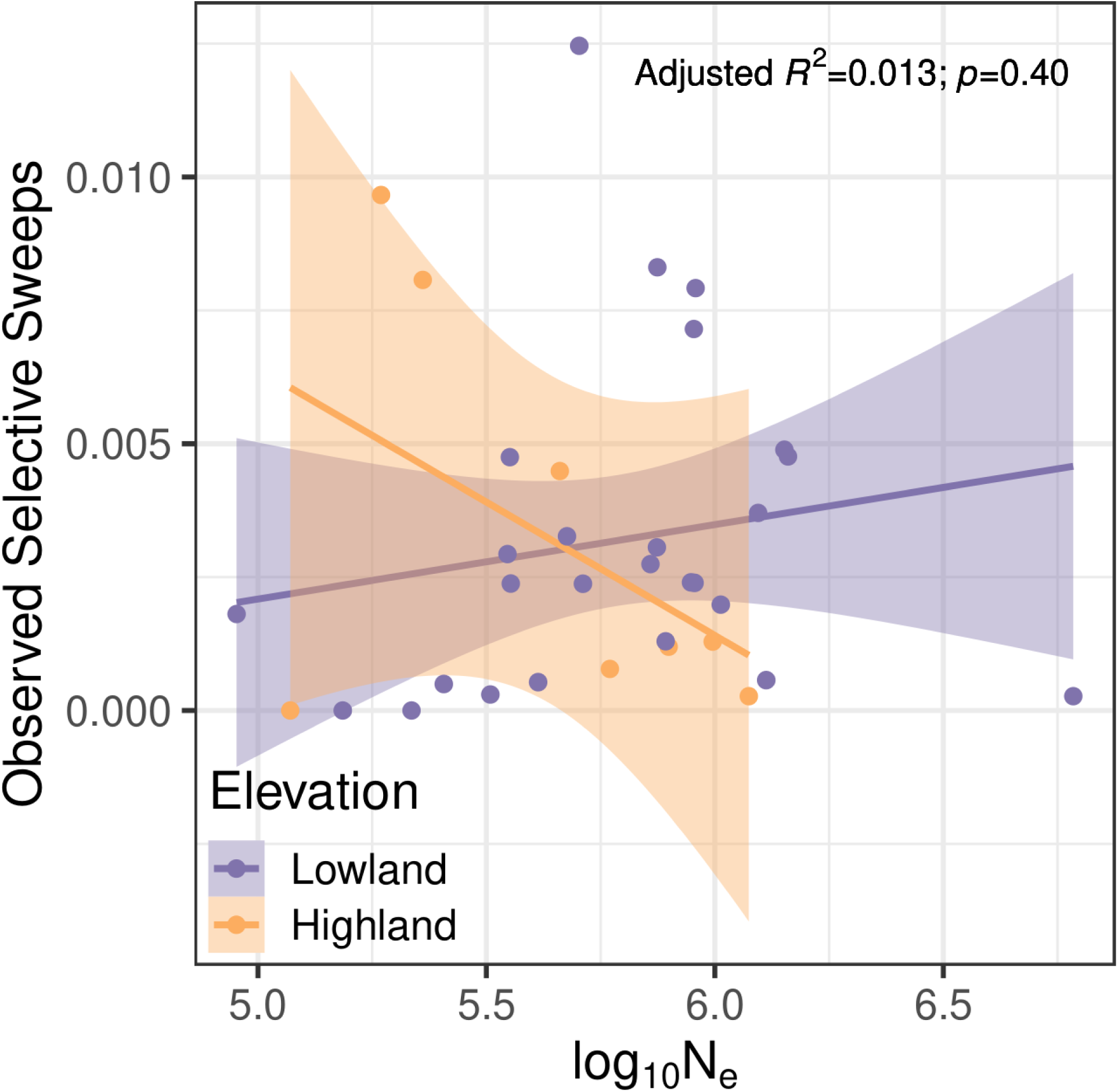
Regression analyses between the number of lineage-specific windows of selection and log_10_ effective population size (log_10_*N*_e_) for individual species. Here, log_10_*N*_e_ interacted with elevational category of lowland or highland. Species from the Andean highlands are shown in orange, and lowland Amazonian species are shown in purple. This model was not significant (*p*=0.40).

**Figure S6.**
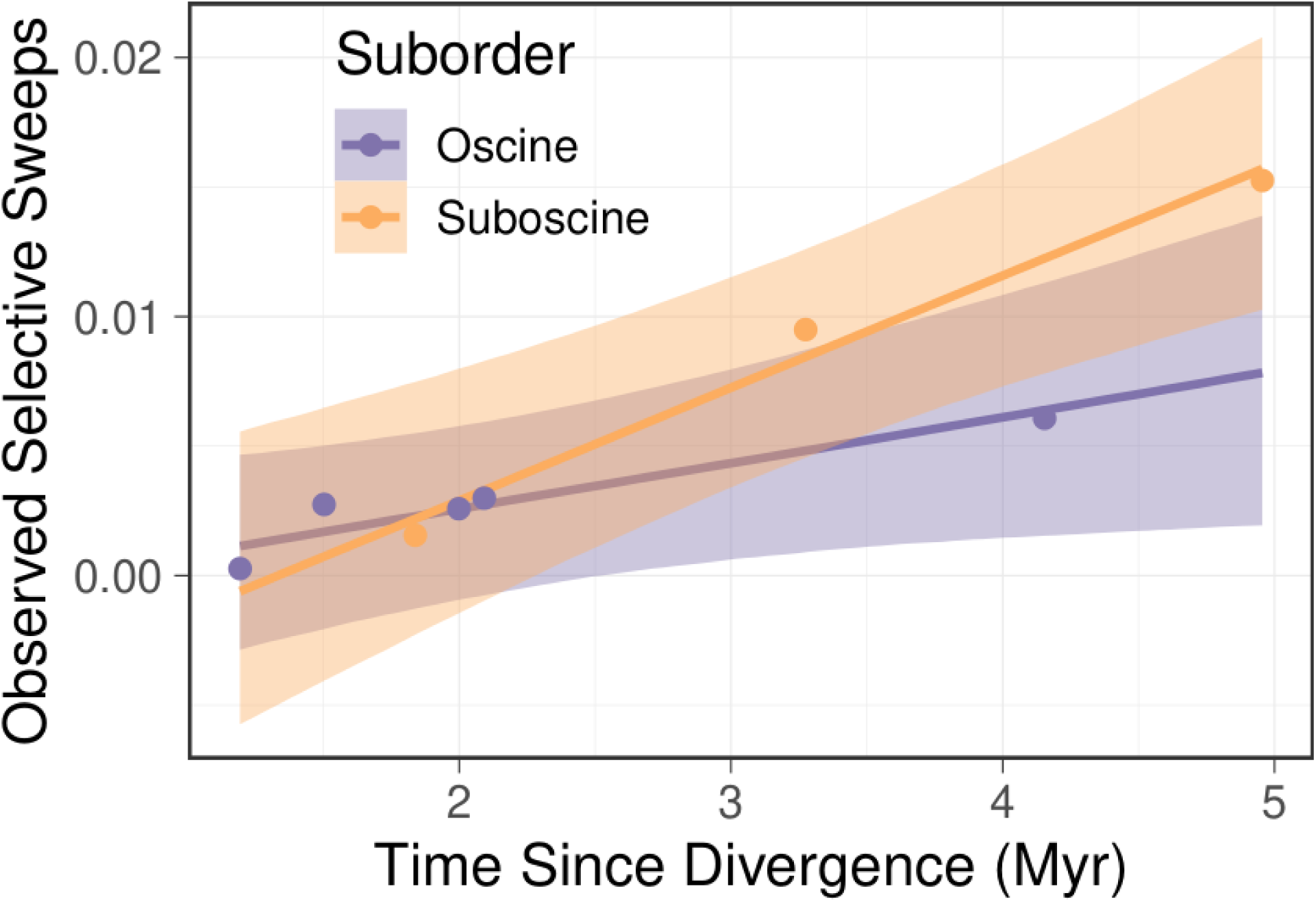
Number of observed selective sweeps accumulated between species pairs versus their divergence times, calculated from the nuclear genome using demographic modeling for only elevationally differentiated species pairs. Oscine species pairs are shown in purple, and suboscine pairs are shown in orange. Lines show model fits with phylogenetic relatedness accounted for. Shading shows the 95% credible intervals.

**Figure S7.**
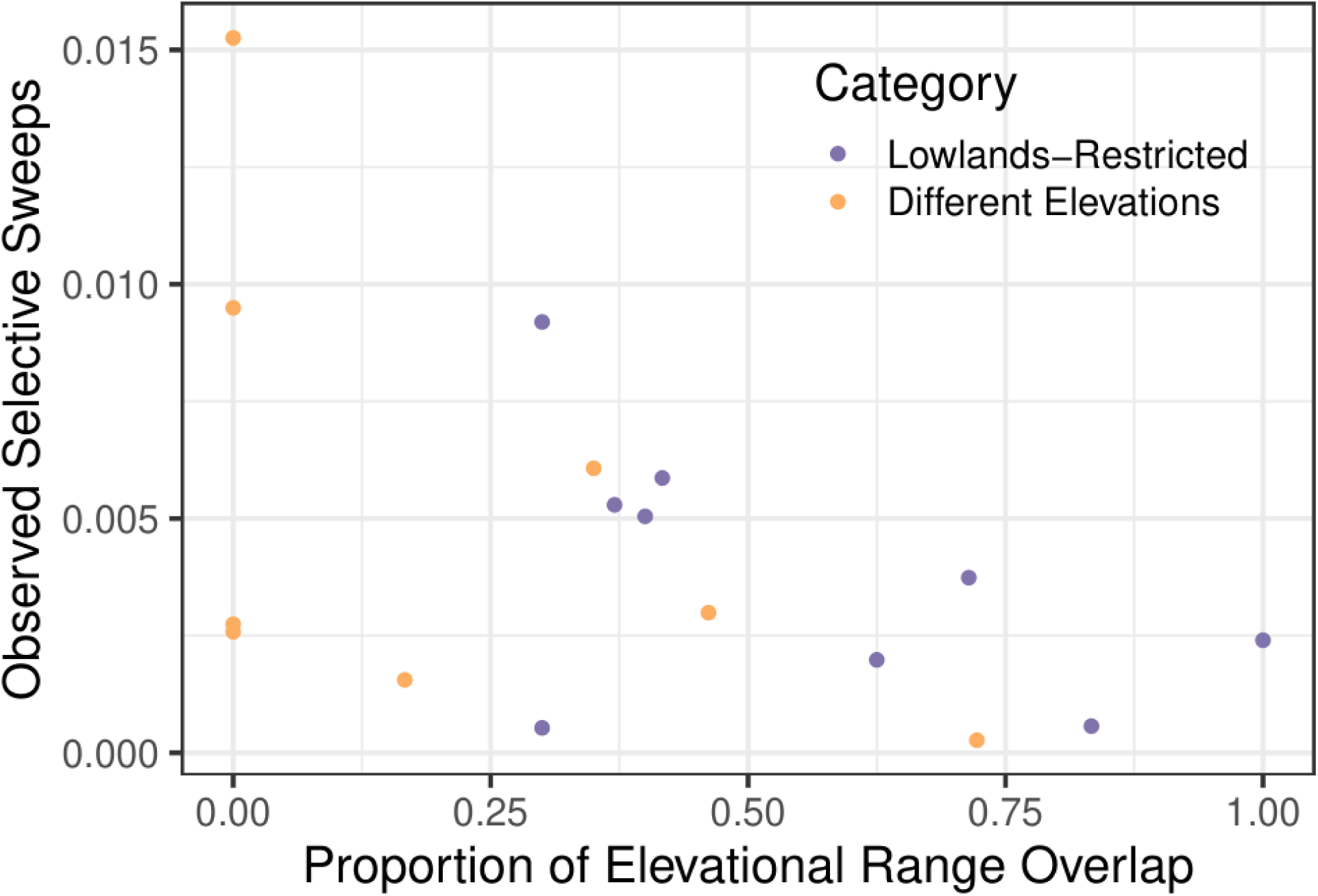
Proportion of elevational range overlap for each sister species pair, within lowlands-restricted or different elevation categories, against observed selective sweeps.

